# Amino acid residues involved in inhibition of host gene expression by influenza A/Brevig Mission/1/1918 PA-X

**DOI:** 10.1101/2021.04.23.441102

**Authors:** Kevin Chiem, Luis Martinez-Sobrido, Aitor Nogales, Marta L. DeDiego

## Abstract

Influenza A virus (IAV) PA-X protein is a virulence factor that selectively degrades host mRNAs leading to protein shutoff. This function modulates host inflammation, antiviral responses, cell apoptosis, and pathogenesis. In this work we describe a novel approach based on the use of bacteria and a plasmid encoding the PA-X gene under the control of the bacteriophage T7 promoter to identify amino acid residues important for A/Brevig Mission/1/1918 H1N1 PA-X’s shutoff activity. Using this system, we have identified PA-X mutants encoding single or double amino acid changes, which diminish its host shutoff activity, as well as its ability to counteract interferon responses upon viral infection. This novel bacteria-based approach could be used for the identification of viral proteins that inhibit host gene expression as well as the amino acid residues responsible for inhibition of host gene expression.

## 1. Introduction

Influenza A viruses (IAV) are single-stranded, negative-sense, segmented viruses belonging to the *Orthomyxoviridae* family, and cause influenza disease [1,2]. IAV are classified into subtypes based on the antigenic properties of the two viral glycoproteins inducing high levels of neutralizing antibodies, the hemagglutinin (HA) and the neuraminidase (NA). Nowadays, 18 different HA subtypes (H1 to H18) and 11 different NA subtypes (N1 to N11) have been described [1–3]. All IAV subtypes (except of H17N10 and H18N11, identified in bats) infect wild aquatic birds, which are considered their natural reservoirs [1–3]. In addition, IAVs are able to infect a wide range of avian and mammalian species, including humans, bats, pigs, dogs, cats, and horses [2–5].

In the last centuries, different IAV subtypes have been responsible of influenza pandemics, including 1889 (H2N2), 1918 (H1N1), 1957 (H2N2), 1968 (H3N2), 1977 (H1N1), and 2009 (H1N1) [6]. These IAV pandemics have caused human deaths varying from ∼2.5% (1918 H1N1) to < 0.1% in other pandemics [7]. The worst influenza pandemic recorded in history was the 1918 “Spanish flu”, responsible for a death toll estimated to be between 20-50 million worldwide [7,8]. Although most experts believe that we will face another influenza pandemic, it is impossible to predict when and where it will originate, or how severe it will be. Therefore, it is important to determine and characterize virulence factors associated to IAV and develop countermeasures against IAV infections.

IAV segment 3 encodes a single, unspliced mRNA which encodes both the polymerase acidic (PA) and the PA-X proteins [9]. The N-terminal domain of PA contains an RNA-endonuclease domain that cleaves capped RNA fragments from cellular pre-mRNAs to provide primers for viral transcription [10,11]. PA-X is translated as a +1 frameshifting from the viral PA mRNA [9], and shares the same first 191 N-terminal amino acids with PA protein, including the endonuclease domain [9]. Influenza PA-X selectively degrades RNAs transcribed by host RNA polymerase II (Pol II), but not other polymerases [12]. mRNA degradation mediated by PA-X leads to cellular gene expression repression and impairment of host antiviral responses [9]. In spite of PA and PA-X sharing the N-terminal endonuclease domain, PA-X shows higher shutoff activity than PA, indicating that the C-terminal PA-X-specific region is also important for this function [10,12–15]. Most of the human IAV, including 1918 pandemic strains, encode a 252 amino acid-length PA-X, with 61 C-terminal amino acids that result from the frameshift and are unique to PA-X [16]. Nevertheless, some IAVs, including the 2009 human pandemic H1N1 (pH1N1), canine and some swine influenza viruses encode a 232 amino acids-length PA-X because of an stop codon at position 42 of the X open reading frame (ORF)[9,17].

Notably, PA-X modulates host inflammation, antiviral responses, cell apoptosis, T-lymphocyte-signaling pathways, and virulence [9] in a strain-specific way [18]. Loss of PA-X expression increased viral replication, host inflammatory response, and virulence in both 2009 pH1N1-and H5N1-infected mice [19,20], as well as in H5N1-infected ducks and chickens [21]. In addition, PA mRNA and protein synthesis was upregulated in cells infected with PA-X-deficient pH1N1 and H5N1 viruses [20]. For 1918 influenza virus, PA-X decreases virulence in a mouse infection model. In addition, loss of PA-X expression increases inflammatory, apoptotic ad T lymphocyte-signaling pathways [9]. In contrast, absence of PA-X in H9N2 IAV decreased viral pathogenicity in mice, reduced progeny virus production, and dampened proinflammatory cytokine and chemokine response [22], suggesting different effects of PA-X protein on virus replication, induction of innate immune responses and pathogenicity depending on the IAV subtype or even strain. Notably, pH1N1, H5N1, and H9N2 IAVs encoding the 252-amino acid length PA-X replicated more efficiently and were more virulent in mice than viruses encoding a truncated 232 amino acid-length PA-X protein [23,24].

Many residues have been shown to be relevant for PA-X’s endonuclease activity, including the catalytic residue K134 and the bivalent cation-binding residue D108 [9,10]. Moreover, six basic amino acid residues (R195, K198, R199, K202, K203, and K206) located at the C-terminal PA-X-specific region, play a key role in PA-X’s ability to dampen cellular host gene expression [25–27]. R195K, K206R, and P210L substitutions conferred significantly increased replication and pathogenicity to H9N2 IAV in mice and ferrets [26]. Amino acid substitutions F4S, F9L, Y24S, D27G, C39Y, C45W, A87V, I94N, L106P/S184I, P107S, D108E, D108N, E119N, I120F, T123I, R124S, R125K, H146Y, E154K, E154A, D160Y, L163R, R168M, and I171M decrease A/WSN/33 H1N1 (WSN) PA-X’s inhibition of host gene expression, as shown using a yeast screening [28]. These data highlight that many amino acid changes can affect PA-X’s activity, and some of them could be strain-dependent. Although residues important for PA-X shutoff activity have been identified using the WSN strain, amino acid residues involved in the ability of 1918 PA-X to inhibit host gene expression have not been yet identified. Moreover, differences or similarities among IAV strains (e.g. WSN and 1918 H1N1 strains) will help to understand the mechanisms of PA-X’s inhibition of host gene expression, similarities and/or differences between amino acid residues involved in host cellular shutoff, and to identify new antivirals targeting key residues in PA-X proteins involved in inhibition of host gene expression.

In this work, we used an innovative bacterial-based approach to identify amino acid residues important for 1918 PA-X’s ability to induce cellular shutoff. This approach is based on the ability of PA-X protein to inhibit bacteria host gene expression and therefore growth. Notably, we have identified novel amino acid residues important for the shutoff activity of 1918 PA-X. This approach could be used to identify other viral proteins inhibiting host gene expression as well as the amino acid residues responsible for their shutoff activity, including other PA-X proteins encoded by different IAV strains.

## 2. Materials and Methods

### 2.1. Cell lines

Human embryonic kidney (HEK) 293T cells (American Type Culture Collection, ATCC, CRL-11268) were cultured and maintained in Dulbecco’s modified Eagle’s medium (DMEM) supplemented with 10% fetal bovine serum (FBS) and 1% PSG (penicillin, 100 units/ml; streptomycin 100 µg/ml; L-glutamine, 2 mM) at 37°C with 5% CO_2_.

### 2.2. Bacterial screening assay and plasmids

The pGBKT7 plasmid encoding 1918 H1N1 wild-type (WT) PA-X gene was generated using specific primers and standard molecular biology methods. The pGBKT7-PA-X plasmid was transformed into chemically competent DH5α *E. coli* (ThermoFisher Scientific), and the transformants were plated on Luria broth (LB) agar plates containing 50 µg/ml kanamycin A and incubated for 24 hours at 30°C. An empty pGBKT7 plasmid and a pGBKT7 plasmid encoding the NS1 gene from A/PuertoRico/8/34 H1N1 strain were included in separate transformations as internal controls. Bacterial colonies were selected and grown in LB containing 50 µg/ml kanamycin A at 30°C for 24 hours and plasmid DNA were purified using the E.Z.N.A Plasmid Mini Extraction kit (Omega Bio-tek) based on the manufacturer’s instructions. Mutant PA-X were cloned into the mammalian expression plasmid pCAGGS containing an HA epitope tag at the N-terminus using Smal and XhoI restriction sites [4]. Sequences for all plasmids containing PA-X mutants were confirmed (ACGT Inc.). Primers used for the generation of plasmid constructs and sequence analysis are available upon request.

### 2.3. Host shutoff assays

Host shutoff activity of 1918 H1N1 WT PA-X on host protein synthesis was determined by transiently transfecting HEK293T cells (5 × 10^4^cells/well, 24-well plate format, triplicates) using Lipofectamine 2000 (LPF2000, Invitrogen) with increasing concentrations (0, 1, 10, or 100 ng) of pCAGGS expression plasmids encoding WT 1918 PA-X together with 250 ng of pCAGGS plasmids expressing the green fluorescent protein (GFP) or Gaussia luciferase (Gluc). Empty pCAGGS plasmid was included as an internal control and to normalize the total amount of transfected plasmid DNA. At 24 hours post-transfection (h p.t.), cellular GFP expression and Gluc activity from tissue culture supernatants were evaluated using a fluorescence microscope, or a Biolux Gaussia luciferase assay kit (New England BioLabs) and a GloMax microplate reader (Promega), respectively. A similar protocol was used to evaluate the ability of 1918 PA-X mutants to inhibit host gene expression.

### 2.4. Inhibition of ISRE promoter activation

To evaluate inhibition of Interferon-stimulated response element (ISRE) promoter activation, HEK293T cells (5 × 10^4^cells/well, 96-well plate format, triplicates) were transiently co-transfected using a calcium phosphate mammalian transfection kit (Agilent Technologies), with 10 ng/well of pCAGGS plasmids encoding the WT or mutant 1918 PA-X proteins, or an empty plasmid as a control, together with 20 ng/well of a simian virus 40 (SV40)-driven Renilla luciferase (Rluc) expression plasmid and 50 ng/well of a plasmid expressing Firefly luciferase (Fluc) under the control of the ISRE promoter (pISRE-Fluc) [29]. At 24 h p.t., cells were washed and infected at a multiplicity of infection (MOI) of 3, with the Sendai virus (SeV), Cantell strain, for ISRE promoter activation [29]. At 21 hours post-infection (h p.i.), cells were lysed using passive lysis buffer (Promega). Luciferase expression in the cell lysates was determined using a dual-luciferase kit (Promega) according to the manufacturer’s guidelines. Measurements were recorded with a microplate reader (Apliskan, Thermo Scientific).

### 2.5. Protein gel electrophoresis and Western blot analysis

HEK293T cells from the host shutoff assays above were harvested and lysed in passive lysis buffer (Promega) for 20 min, and proteins from whole cell lysates were separated by denaturing electrophoresis using 12% SDS-polyacrylamide gels. Next, proteins were transferred onto nitrocellulose membranes (Bio-Rad) using a Bio-Rad Mini Protean II electroblotting apparatus at 100V for 2 h. After transfer, membranes were blocked for 1 h with 5% dried skim milk in 1X PBS containing 0.1% Tween 20 (T-PBS) and incubated with primary anti-HA epitope tag (Sigma, H6908) or anti-GFP (Sigma, GSN149) rabbit polyclonal antibodies (P)Ab overnight at 4°C. Mouse monoclonal (M)Ab against β-actin (Sigma, A1978) was included as a loading control. Secondary horseradish peroxidase (HRP)-conjugated Abs (Sigma) specific against mouse MAbs or rabbit PAbs were used to detect the membrane-bound primary Abs. Proteins were detected by chemiluminescence using SuperSignal West Femto substrate (ThermoFisher Scientific) following the manufacturer’s specifications and imaged using a ChemiDoc Imaging System (Bio-Rad).

### 2.6. Immunofluorescence assay

To assess cellular PA-X expression by indirect immunofluorescence, HEK293T (24-well plate format, 2.5 × 10^5^cells/well, triplicates) were transiently transfected, using LPF2000, with 1 µg of the indicated pCAGGS expression plasmids encoding 1918 PA-X WT or mutants. At 24 h p.t., cells were fixed and permeabilized with 4% (vol/vol) formaldehyde and 0.5% (vol/vol) Triton X-100 (Sigma), respectively, before blocking with 1X PBS containing 2.5% FBS for 1 h. Thereafter, cells were incubated with an anti-HA epitope tag PAb (Sigma, H6908) and then with a FITC-conjugated donkey anti-rabbit secondary Ab (Invitrogen). Cell nuclei were stained with DAPI (4′,6-diamindino-2-phenylindole). Representative images were obtained using an EVOS M5000 Imaging System (ThermoFisher Scientific).

### 2.7. Structure Analysis

Amino acid positions were plotted on the crystal structure of the N-terminal region of pH1N1 PA protein (Accession #AY818132) using the PyMOL molecular graphics system.

### 2.8. Statistical Analysis

Microsoft Excel (Microsoft Corporation) and GraphPad Prism software were used to analyze the data. Microsoft Excel was necessary to perform some of the calculations, and to visualize the raw data. One-way analysis of variance (ANOVA) was performed using GraphPad Prism software.

## 3. Results

### 3.1. Inhibition of host gene expression by A/Brevig Mission/1/1918 PA-X

To confirm whether the PA-X protein from 1918 possesses the ability to inhibit host gene expression in our assay, as previously described [9], HEK293T cells were co-transfected with pCAGGS expression plasmids encoding GFP and Gluc, together with increasing concentrations of a pCAGGS plasmid expressing an N-terminally HA epitope tagged WT PA-X from 1918 strain **(Figure 1**). At 24 h p.t., GFP expression and Gluc activity from the cell culture supernatants were evaluated, showcasing a dose-dependent decline in GFP expression (**Figure 1A**) and Gluc activity (**Figure 1B**), indicative of 1918 PA-X protein-mediated inhibition of cellular gene expression. Whole cell lysates were evaluated by Western blot, in which the PA-X protein was only detectable in cells transfected with 100 ng of plasmid (**Figure 1C**). Conversely, greater amounts of GFP were detected by Western blot with less 1918 PA-X-expressing pCAGGS plasmid (**Figure 1C**), analogous to results observed by fluorescence microscopy (**Figure 1A**). These results support the ability of 1918 PA-X to inhibit host gene expression.

**Figure 1.**
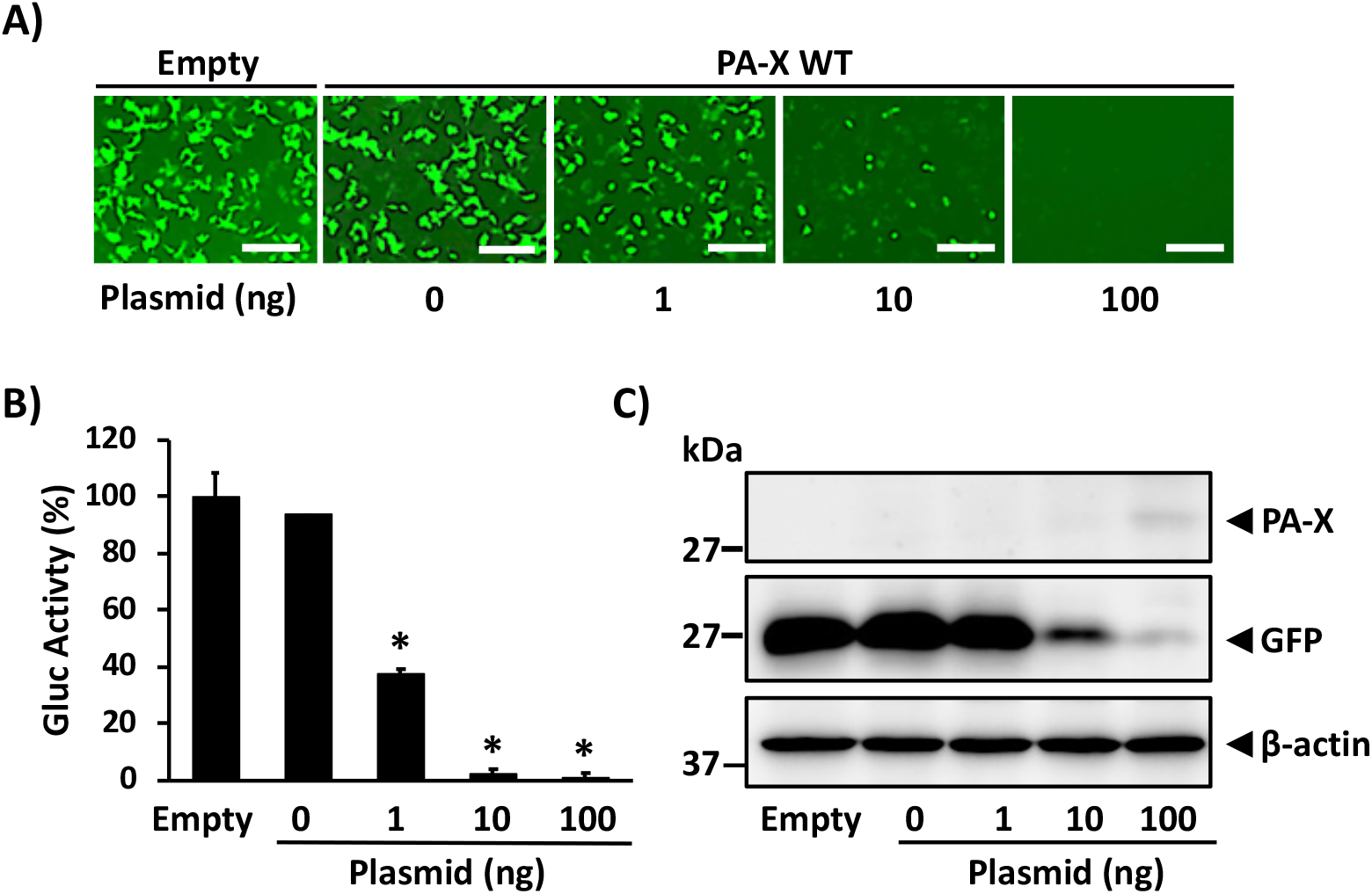
Ability of 1918 PA-X to inhibit host gene expression: HEK293T cells were co-transfected with increasing concentrations (0, 1, 10, or 100 ng/well) of a pCAGGS expression plasmid encoding wild-type 1918 PA-X (PA-X WT) fused to an HA epitope tag, and 250 ng of pCAGGS-Gluc and -GFP plasmids. Empty plasmid transfected cells were included as control. At 24 h p.t., GFP was observed under a fluorescence microscope (**A**), and Gluc activity was quantified using a GloMax system from cell culture supernatants (**B**). Gluc activity was normalized to empty plasmid transfected cells (100%). *, P < 0.0001; (0 ng of PA-X WT plasmid versus other concentrations) using one-way ANOVA. The presence of PA-X and GFP in cell lysates was analyzed by Western blot using anti-HA or anti-GFP pAbs, respectively (**C**). β-actin was used as a loading control. Results are the means and SDs of triplicates. Data are representative of three separate experiments. Magnification 20X, Scale bar, 100 µm.

### 3.2. Identification of PA-X mutants impaired in its ability to inhibit host gene expression

In order to identify 1918 PA-X amino acid residues responsible for its host shutoff activity, we cloned the 1918 PA-X open reading frame into the pGBKT7 yeast expression plasmid using DH5α bacteria competent cells. Remarkably, an unnaturally low number of bacterial colonies were acquired compared to an experimental approach involving the same pGBKT7 empty plasmid or a pGBKT7 plasmid encoding influenza A/Puerto Rico/8/34 H1N1 NS1 gene. We and others have previously found that proteins cloned under a T7 promoter in plasmids were expressed in bacteria (data not shown) [30]. Therefore, we postulated that low levels of expression of WT 1918 PA-X from the pGBKT7 plasmid was deleterious and/or toxic to bacteria [30], due to the ability of PA-X to inhibit host gene expression, in contrast to A/Puerto Rico/8/34 H1N1 NS1 protein, which does not significantly inhibit host gene expression [29]. In light of this, we hypothesized that the recovered bacterial colonies must contain plasmids encoding mutated forms of the 1918 PA-X that affect its shutoff capabilities. The genetic sequencing of 1918 PA-X-encoded pGBKT7 plasmids extracted from the few recovered bacterial colonies revealed single or double mutations that causes amino acid changes in 1918 PA-X protein, and some early or late stop codons leading to deletions and/or insertions within the 1918 PA-X gene (**Table 1**).

**Table 1.** Amino acid residues in 1918 PA-X identified in the bacterial-based assay.

| Amino acid (aa) change | Number of clones<sup>1</sup> | PMID Reference<sup>2</sup> |
| --- | --- | --- |
| M21V | <u>1</u> |  |
| F76L | <u>2</u> |  |
| L109R | <u>1</u> |  |
| T123A | <u>1</u> | 29331676 |
| R168G | <u>2</u> | 29331676 |
| I79V/Y161L | <u>2</u> |  |
| T98I/P103S | <u>1</u> |  |
| H146Y/L187P | <u>1</u> | H146Y<br>(29331676) |
| 94 aa truncation | <u>1</u> |  |
| 180 aa truncation | <u>1</u> |  |
| No stop, additional 23 aa | <u>1</u> |  |
| 1 nucleotide deletion, early stop | <u>1</u> |  |

<sup>1</sup>Number of clones identified for each mutation in the bacterial-based assay.
<sup>2</sup>If the identified amino acid change has been previously described to be involved in inhibition of host gene expression.

Out of 15 bacterial colonies, all isolated pGBKT7 plasmids encoded 1918 PA-X genes with mutations, five of them containing unique single (M21V, F76L, L109R, T123A, and R168G) mutations and three of them containing double (I79V/Y161C, green; T98I/P103S, blue; and H146Y/L187P, magenta) amino acid mutations (**Figure 2 and Table 1**). All of these identified mutations caused amino acid changes in both the PA-X and PA proteins, which we hypothesize affect the host shutoff activity of PA-X. Notably, the same amino acid changes F76L, R168G, I79V/Y161C were found in plasmids from two independent bacteria colonies (**Table 1**).

**Figure 2.**
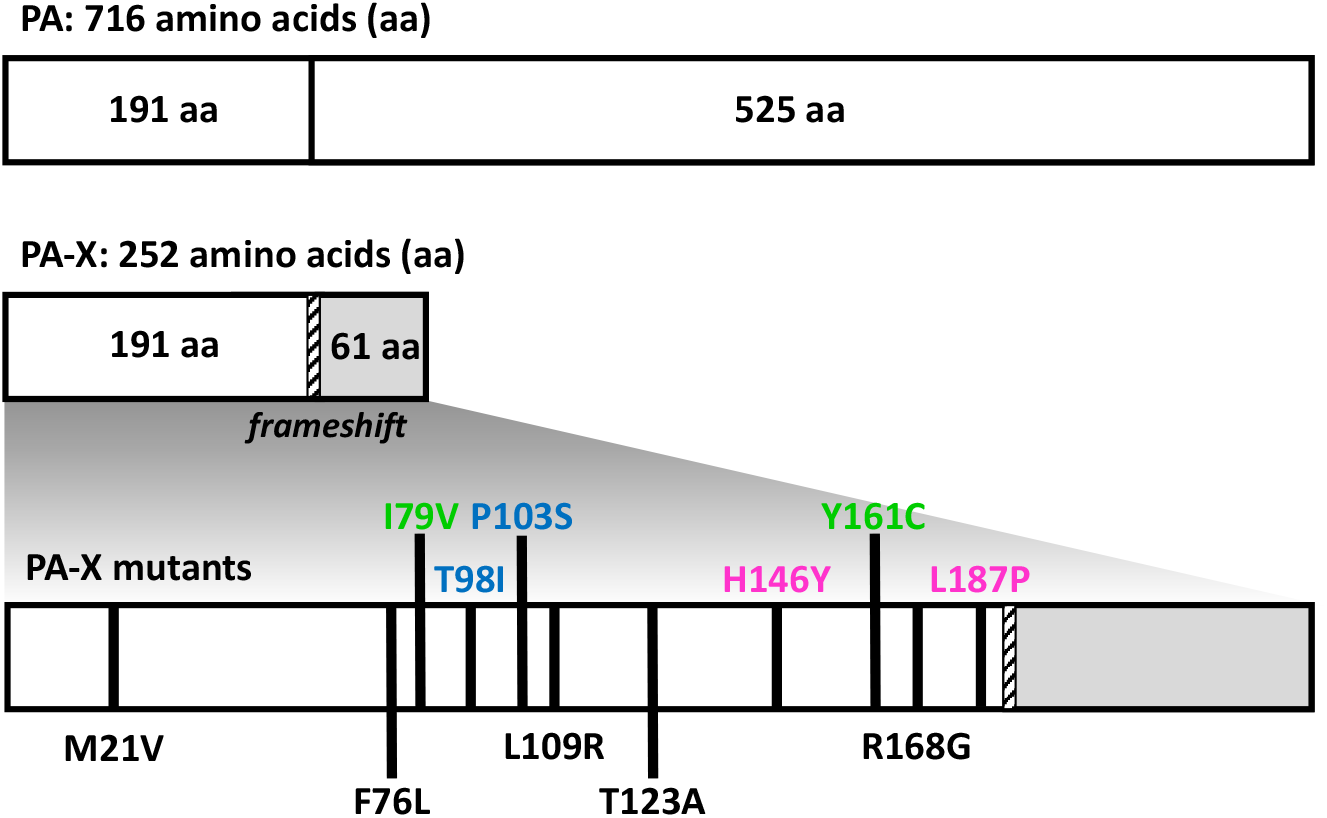
Schematic representation of 1918 PA and PA-X identified mutations: PA-X protein is encoded from the viral PA segment using a +1 frameshift motif (striped bar). Using a bacteria-based assay, amino acid residues in 1918 PA-X involved in host shutoff were identified (black lines). PA-X clones containing a single mutation are indicated in black. PA-X clones containing double mutations are paired by either green, blue, or magenta.

### 3.3. Effect of amino acid changes in 1918 PA-X-mediated shutoff activity

Upon sequencing of pBGKT7 plasmids containing the PA-X gene from the recovered bacteria, several mutations were identified, suggesting their importance for host shutoff activity. To investigate this, we assessed the ability of the identified 1918 PA-X mutants to inhibit host gene expression and compared them to that of the WT 1918 PA-X (**Figure 3**). Each of the 1918 PA-X mutants displayed weakened, albeit varying degrees, inhibition of host gene expression compared to WT 1918 PA-X, as indicated by GFP expression (**Figure 3A**) and Gluc activity (**Figure 3B**). Notably, PA-X mutations L109R, T123A, T98I/P103S and H146Y/L187P had a greater impact on PA-X’s ability to suppress host gene expression (**Figures 3A and 3B**). These results were further validated by Western blot analysis (**Figure 3C**) because of the ability of 1918 PA-X to inhibit its own expression when expressed under the control of a RNA polymerase II promoter [12]. As expected, all 1918 PA-X mutants were expressed to higher levels than 1918 WT PA-X (**Figure 3C**). Similarly, GFP and Gluc expression levels were higher in cells transfected with 1918 PA-X mutants (**Figures 3A and 3B**), predominantly L109R, T123A, T98I/P103S and H146Y/L187P, whose inhibitory activity is greatly affected. Surprisingly, PA-X containing a T123A amino acid change showed the highest expression levels by Western blot (**Figure 3C**). Although the reason(s) for these differences were not directly addressed, it could be due to protein stability. Recently, it has been described that the half-life of PA-X ranges from 30 min to 3.5 h depending on the IAV strain [31]. Notably, sequences in the C- and N-terminal domains were important for regulating PA-X half-life [31].

**Figure 3.**
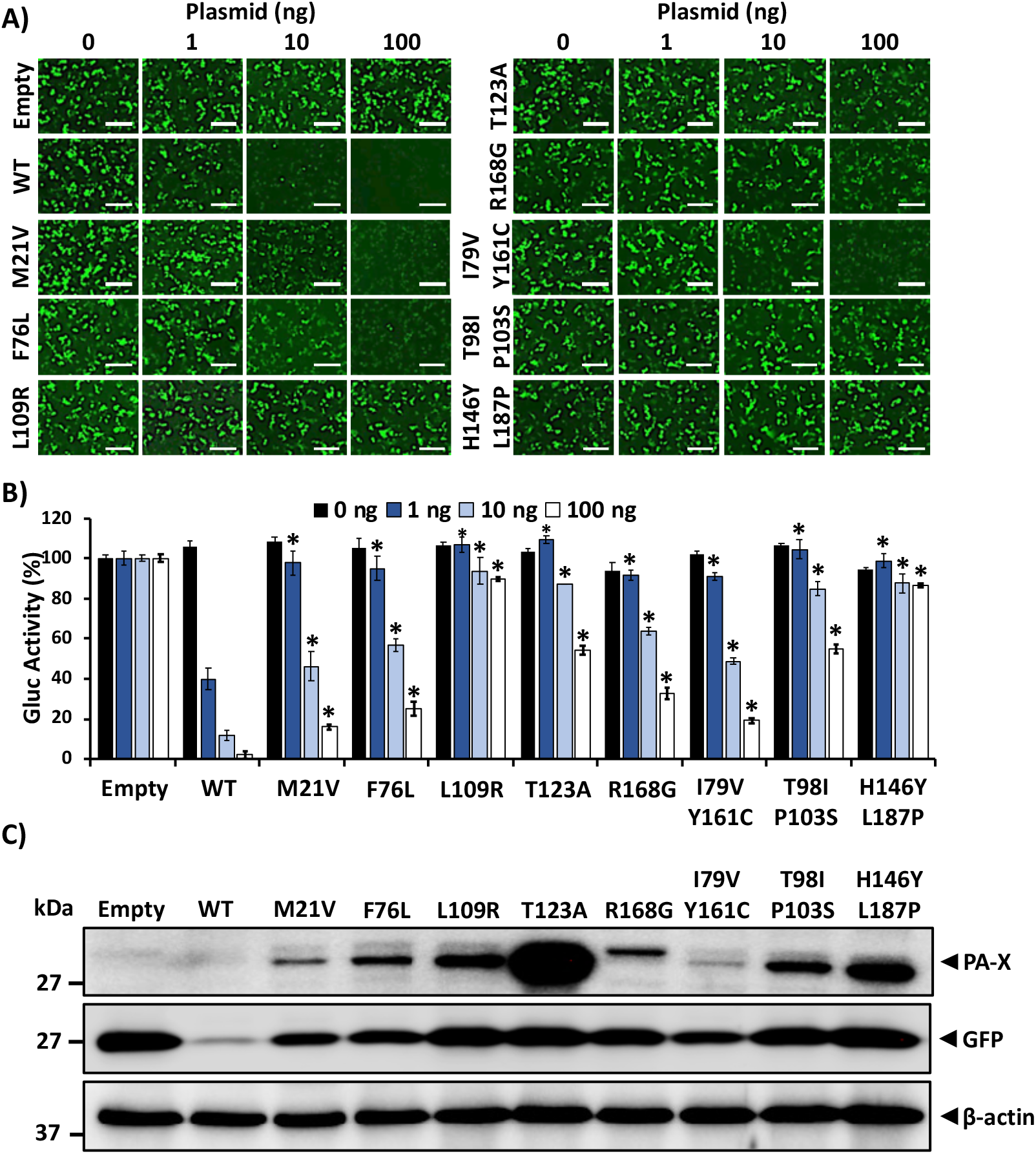
Ability of 1918 PA-X mutants to inhibit host gene expression: HEK293T cells were co-transfected with increasing concentrations (0, 1, 10, or 100 ng/well) of pCAGGS expression plasmids encoding the identified 1918 PA-X mutants and 250 ng of pCAGGS-Gluc and -GFP plasmids. Empty and 1918 WT PA-X pCAGGS plasmids were included as controls. At 24 h p.t., GFP was observed under a fluorescence microscope (**A**), and Gluc activity was quantified from cell culture supernatants (**B**). Gluc activity was normalized to the empty plasmid control transfected cells (100%). *, P < 0.0001; (PA-X WT versus PA-X mutants for each amount of plasmid) using one-way ANOVA. PA-X and GFP expression levels in cell lysates were analyzed by Western blot using anti-HA or anti-GFP pABs, respectively (**C**). β-actin was used as a loading control. Results are the means and SDs of triplicates. Data are representative of three separate experiments. Magnification 20X, Scale bar, 100 µm.

Next, we evaluated the expression of 1918 PA-X mutants by immunofluorescence in HEK293T cells transfected with pCAGGS plasmids expressing each of the identified 1918 PA-X mutants (**Figure 4**). Empty and 1918 WT PA-X-expressing pCAGGS plasmids were included as controls. Unsurprisingly, 1918 WT PA-X was not detected in transfected cells by immunofluorescence because its ability to inhibit host gene expression, including its own expression (**Figure 4**) [12,25,27,32–36]. However, 1918 PA-X containing F76L, L109R, T123A, R168G, T98I/P103S and H146Y/L187P mutations were detectable under immunofluorescence because these 1918 PA-X mutants were affected in its shutoff capabilities (**Figure 3**). These results demonstrate that the identified 1918 PA-X mutants are affected in their abilities to induce cellular shutoff.

**Figure 4.**
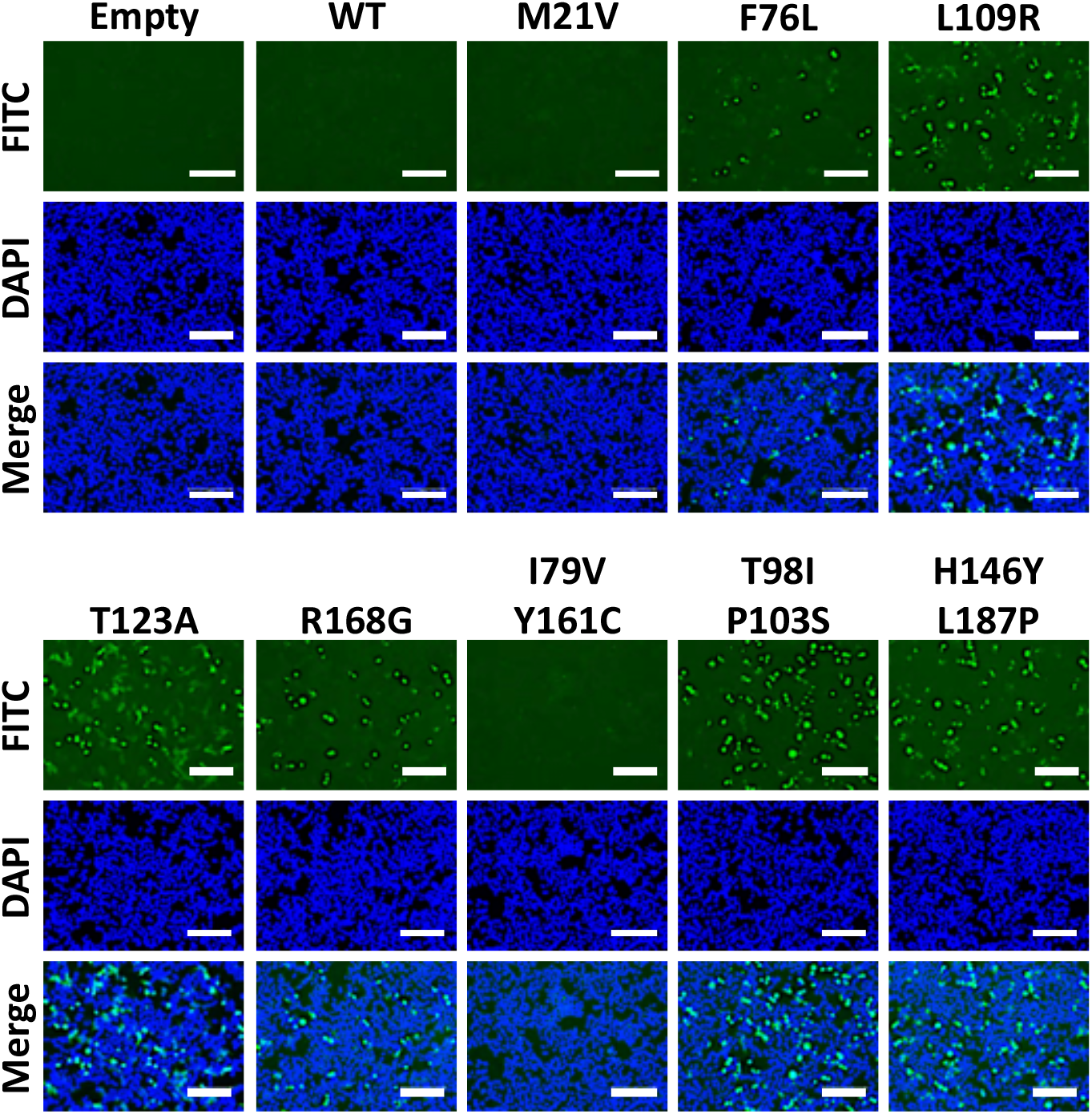
Expression of 1918 PA-X WT and mutants by immunofluorescence: HEK293T cells were transfected with 1 µg of pCAGGS 1918 PA-X HA-tagged WT or the indicated mutants. Empty pCAGGS transfected HEK293T cells were included as internal control. At 24 h p.t., cells were fixed, permeabilized and immunostained with a pAb against the HA epitope tag (green). DAPI was used to stain the nucleus of transfected cells (blue). Magnification 20X, Scale bar = 100 µm.

In the case of the double I79V/Y161C, T98I/P103S and H146Y/L187P mutants, two amino acid changes in 1918 PA-X protein could be involved in their impaired inhibition of host gene expression (**Figure 2** and **Table 1**). To determine whether one specific residue, or both, were critical for 1918 PA-X mutants ability to inhibit host gene expression, pCAGGS-PA-X expressing plasmids encoding each individual amino acid residue were constructed and tested in the cell-based host gene inhibition assay by assessing GFP expression (**Figure 5A**) and Gluc activity from tissue culture supernatants (**Figure 5B**) of transfected HEK293T cells. Expectedly, the greatest amount of GFP signal and Gluc activity was observed for the double mutants (I79V/Y161C, T98I/P103S and H146Y/L187P) transfected cells. However, single mutants displayed decreased GFP and Gluc activity, mainly I79V, Y161C, T98I, and H146Y alone. Yet, L187P and P103S maintained high GFP and Gluc activity, almost to the levels of the double mutant T98I/P103S and H146Y/L187P. In Western blot analysis, we found detectable levels of PA-X in all of the double mutants, and a weaker signal in single mutant P103S and L187P (**Figure 5C**), which were parallel to the expression levels of GFP (**Figure 5A**) and Gluc activity (**Figure 5B**).

**Figure 5.**
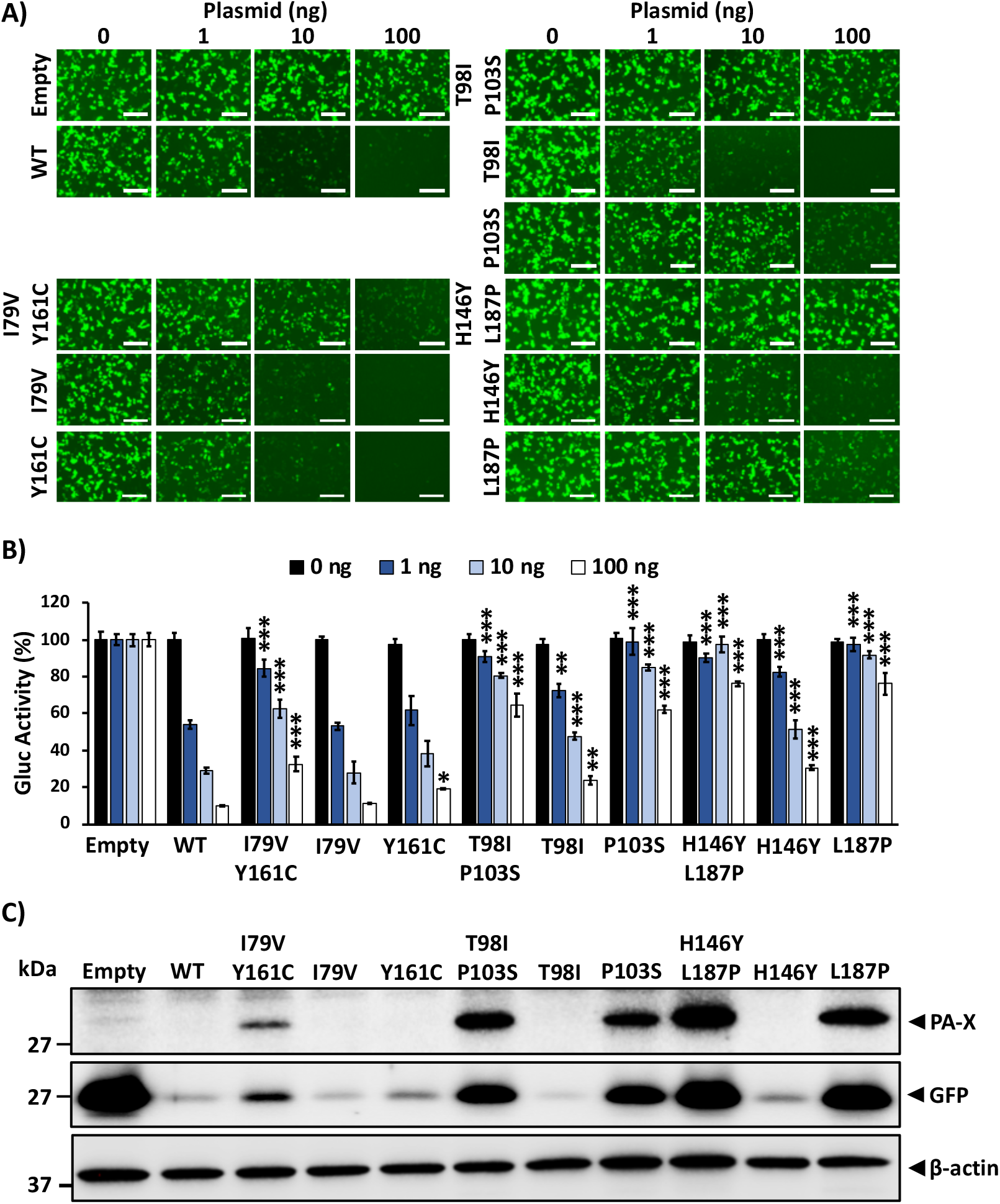
Amino acid mutations of 1918 PA-X double mutants involve in inhibition of host gene expression: HEK293T cells were co-transfected with increasing concentrations (0, 1, 10, or 100 ng/well) of pCAGGS expression plasmids encoding 1918 PA-X single or double mutants fused to an HA epitope tag, and 250 ng of pCAGGS-Gluc and -GFP plasmids. Empty and 1918 WT pCAGGS plasmids were included as control. At 24 h p.t., GFP was observed under a fluorescence microscope (**A**), and Gluc activity was quantified using from cell culture supernatants (**B**). Gluc activity was normalized to the empty plasmid control transfected cells (100%). *, P < 0.05; **, P < 0.0005; ***, P < 0.0001; (PA-X WT versus PA-X mutants for each amount of plasmid) using one-way ANOVA. PA-X and GFP expression levels in cell lysates were analyzed by Western blot using anti-HA or anti-GFP pAbs, respectively (**C**). β-actin was used as a loading control. Results are the means and SDs of triplicates. Data are representative of three separate experiments. Magnification 20X, Scale bar, 100 µm.

Lastly, 1918 PA-X double and single mutants were evaluated by immunofluorescence, which displayed no signal for WT, I79C/Y161C, or its single mutants I79V or Y161C (**Figure 6**). However, double mutants T98I/P103S andH146Y/L187P, along with their single mutant P103S and L187P were detectable in these immunofluorescence assay (**Figure 6**). Overall, these results demonstrate that the combination of I79V and Y161C are both important for affecting PA-X ability to inhibit host gene expression. However, in the case of double mutant T98I/P103S and H146Y/L187P, P103S and L187P are mainly responsible for the lack of host shutoff activity of 1918 PA-X, although this inhibition is enhanced by T98I and H146Y, respectively.

**Figure 6.**
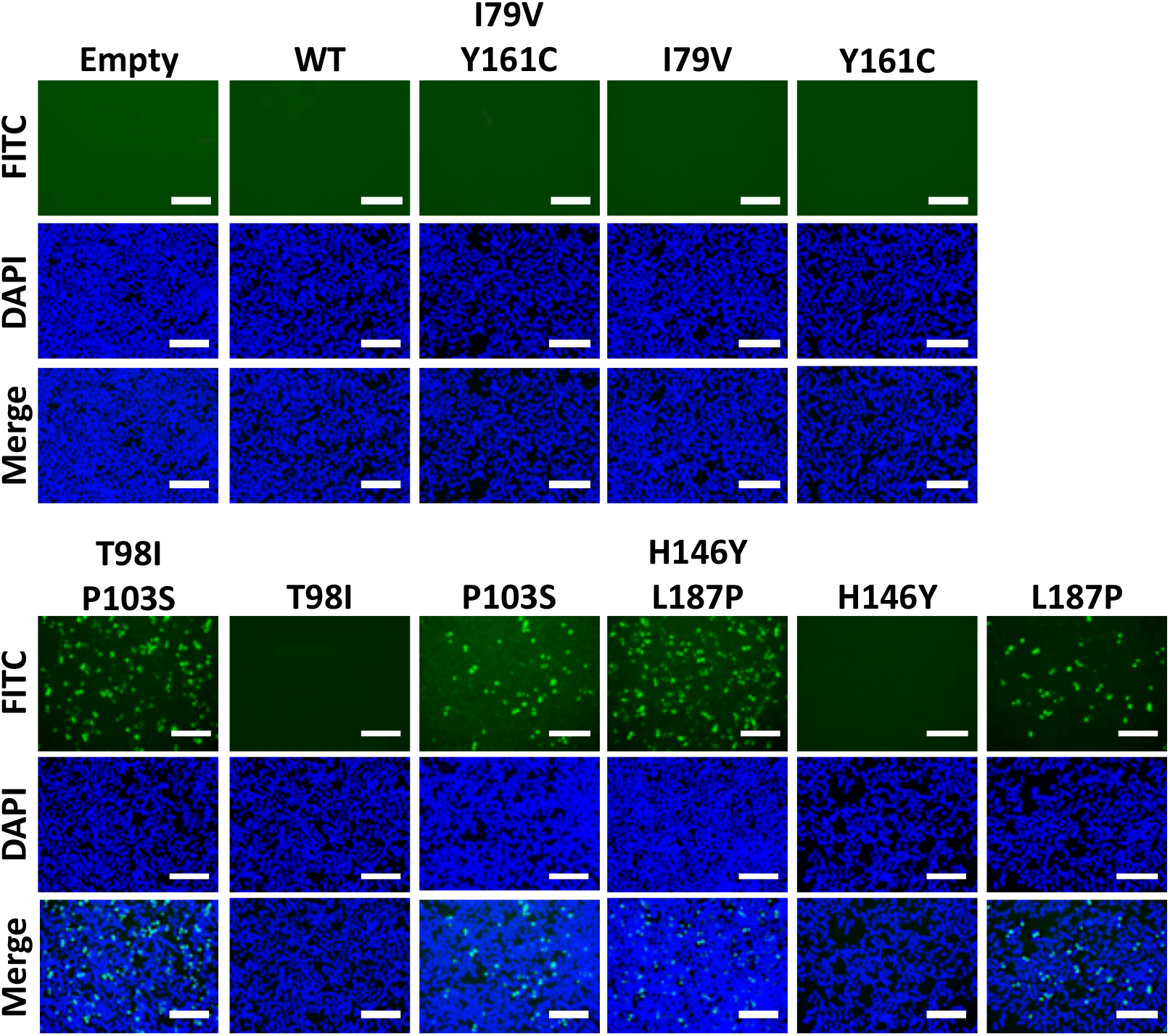
Expression of 1918 PA-X double and single mutants by immunofluorescence: HEK293T cells (24-well plate format, 5 x 104 cells/well) were transfected with 1 µg of pCAGGS 1918 PA-X HA-tagged double or single mutant plasmids. Empty and 1918 PA-X WT pCAGGS expression plasmids were included as internal controls. At 24 h p.t., cells were fixed, permeabilized and immunostained with a pAb against the HA epitope tag (green). DAPI was used to stain the nucleus of transfected cells (blue). Magnification 20X, Scale bar = 100 µm.

### 3.4. Effect of identified PA-X amino acid changes on inhibition of IFN responses

IAV PA-X protein has been shown to counteract innate immune responses due to its effect on host cellular expression [32,33,37]. To analyze the effect of the amino acid changes identified in modulating innate immune responses, we used a well-established virus-based assay. To this end, HEK293T cells were co-transfected with pCAGGS plasmids expressing 1918 PA-X WT or mutants, together with a plasmid expressing Fluc under the control of an ISRE promoter (**Figure 7A**) and a plasmid constitutively expressing Rluc (SV40-Rluc) to evaluate shutoff activity (**Figure 7B**). At 24 h p.t, cells were mock-infected or infected (MOI of 3) with SeV and reporter expression levels were evaluated at 21 h p.i by measuring expression levels of Fluc (**Figure 7A**) and Rluc (**Figure 7B**). In cells transfected with empty plasmid, SeV infection induced high levels of Fluc expression driven by the ISRE promoter (**Figure 7A**). However, activation of ISRE promoter in cells transfected with 1918 PA-X proteins was significantly reduced (**Figure 7A**). These data is consistent with previous data showing that PA-X protein efficiently counteracts IFN responses [13,16,33,38]. Interestingly, 1918 PA-X mutants inhibiting ISRE promoter activation to higher extend (L109R, T123A, R168G, T98I/P103S and H146Y/L187P) were the ones showing higher shutoff activity. In addition, shutoff activity for 1918 PA-X WT or the identified mutants, measured by Rluc levels (**Figure 7B)**, showed similar results compared to those previously described (**Figure 3**), indicating a positive correlation between 1918 PA-X’s ability to inhibit host gene expression and blocking IFN responses.

**Figure 7.**
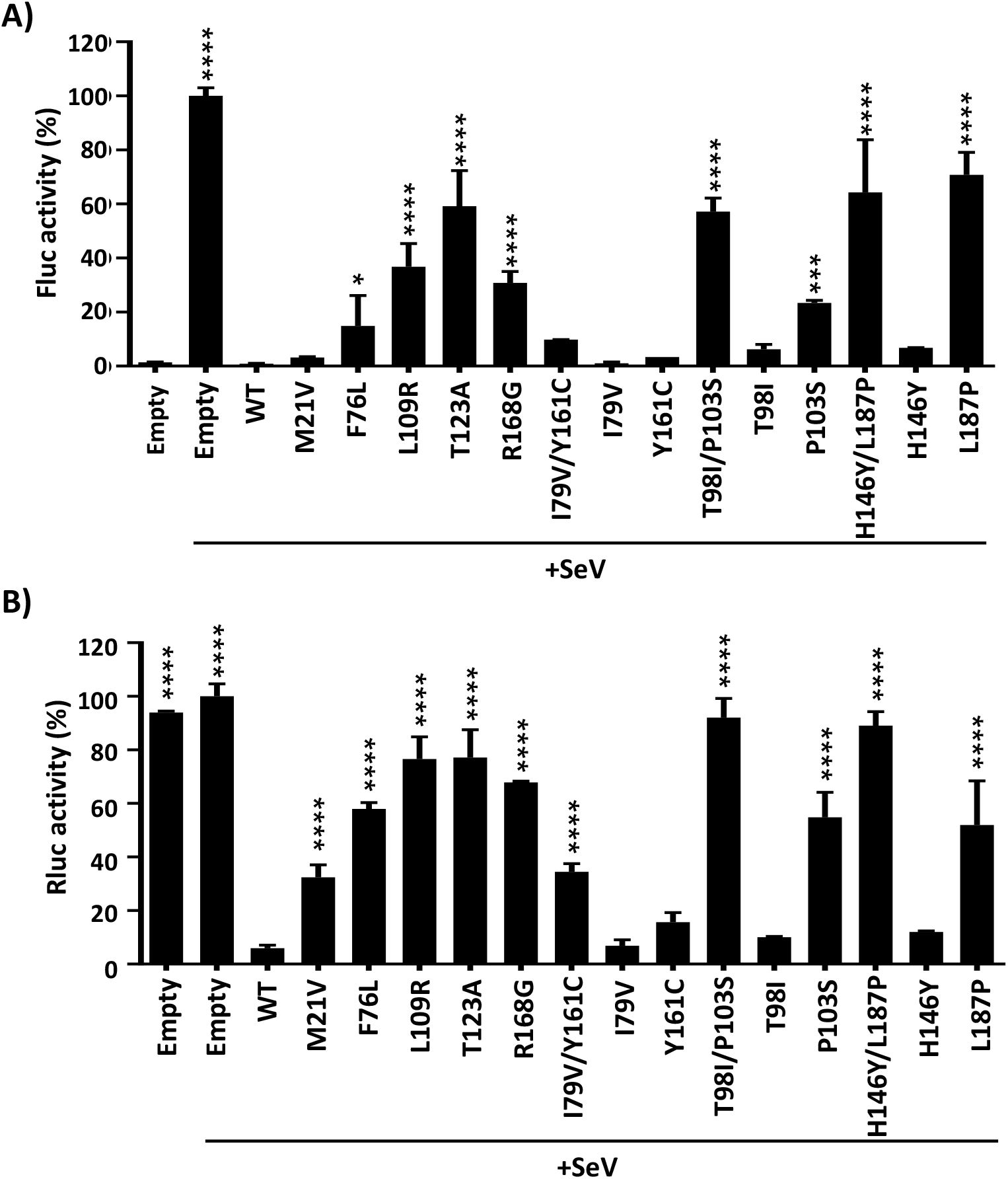
Effect of 1918 PA-X mutants on IFN-induced responses upon SeV infection. (**A** and **B**) HEK293T cells were transiently co-transfected with the indicated HA-tagged WT or mutant 1918 PA-X pCAGGS expression plasmids, together with plasmids expressing Fluc under the control of an ISRE promoter (**A**), or under an SV40 promoter (**B**). Empty pCAGGS plasmid was included as internal control. At 24 h p.t, cells were mock-infected or infected (MOI of 3) with SeV (+SeV) to induce ISRE promoter activation. At 21 h.p.i, cell lysates were prepared for reporter gene expression. Fluc (**A**) and Rluc (**B**) expression levels were measured by luminescence. Activity of Fluc and Rluc were normalized to the empty plasmid control (+SeV) transfected cells (100%). Results are the means and SDs of triplicates. Data are representative of three independent experiments. *, P < 0.05; **, P < 0.005; ***, P < 0.0005; ****, P < 0.0001; (1918 PA-X WT versus 1918 PA-X mutants for each amount of plasmid) using one-way ANOVA.

### 3.5. PA-X amino acid conservation

Sequences available in the Influenza Research Database (https://www.fludb.org/) were aligned to study the conservation of the amino acid residues identified in our bacteria-based screening in the PA-X proteins of H1N1 IAV. Interestingly, the residues found in the PA-X WT proteins from H1N1 IAV are highly conserved and more than 99.7% of the H1N1 IAV isolates encode the WT sequence (**Table 2**). These results suggest that amino acid changes affecting the 1918 PA-X’s ability to inhibit host gene expression are not present in viruses circulating globally, most likely due to a decrease in viral fitness. However, as the identified mutations in PA-X also affect PA protein, we cannot rule out that the negative effect of these mutations on the viruses is due to the PA-X protein, PA protein, or both.

**Table 2.** Amino acid conservation in H1N1 PA-X protein.

| Amino acid change | Amino acid (%) in human sequences (n = 15445)<sup>1</sup> |
| --- | --- |
| M21V | **<u>M (99.99)</u>**<br>I (0.01) |
| F76L | **<u>F (100)</u>** |
| I79V | **<u>I (99.98)</u>**<br>V (0.02) |
| T98I | **<u>T (99.93)</u>**<br>A (0.07) |
| P103S | **<u>P (99.98)</u>**<br>S (0.0065)<br>H (0.0065)<br>L (0.0065) |
| L109R | **<u>L (99.99)</u>**<br>F (0.01) |
| T123A | **<u>T (100)</u>** |
| H146Y | **<u>H (100)</u>** |
| Y161L | **<u>Y(99.74)</u>**<br>F (0.22)<br>H (0.03) |
| R168G | **<u>R (99.99)</u>**<br>T (0.01) |
| L187P | **<u>L (99.92)</u>**<br>I (0.08) |

<sup>1</sup> All H1N1 PA-X sequences available in the Influenza Research database (https://www.fludb.org) from viruses isolated in humans since 1918 were analyzed (N=15445 sequences). Amino acid residues in A/BrevigMission/1918 H1N1 PA-X WT are bold and underlined. Frequency of IAV PA-X proteins containing the indicated residue are shown.

## 4. Discussion

In this work, we show that 1918 PA-X protein, similarly to the PA-X of other IAV subtypes and strains [12,25,27,32–35] is able to impair host protein expression in cells (**Figure 1**). Furthermore, we used a novel approach based in the use of plasmids encoding the 1918 PA-X gene under the control of the T7 bacteriophage promoter to identify amino acid residues important for this shutoff activity (**Figure 2**). Our hypothesis was that 1918 PA-X expression, which was likely driven by the leaking T7 promoter in bacteria [30], was deleterious and/or toxic for bacteria growth, and therefore, bacteria were forced to introduce mutations affecting 1918 PA-X translation or its ability to inhibit host gene expression. In accordance to this hypothesis, we found insertion and/or deletions of nucleotides leading to early or late stop codons within 1918 PA-X gene (**Table 1**). In addition, we also identified 5 different plasmids encoding single amino acid changes in 1918 PA-X gene, and 3 different plasmids encoding two amino acid changes each in 1918 PA-X (**Figure 2 and Table 1**), supporting the hypothesis that bacteria were under selective pressure to introduce changes in the plasmids to abolish the ability of PA-X to inhibit host gene expression. Importantly, all these amino acid changes affected the ability of 1918 PA-X to induce cellular shutoff, measured by GFP and Gluc levels (**Figures 3A, 3B, 5A and 5B**). Furthermore, the identified amino acid changes affected its own expression, as measured by Western blot (**Figures 3C and 5C**) and by immunofluorescence (**Figures 4 and 6**). This was expected as in these experiments genes are transcribed by the RNA polymerase II promoter, and in this scenario 1918 PA-X inhibits its own expression, as previously described [12,27]. It has been shown that amino acid residues located at the C-terminal unique region of PA-X are highly relevant for its shutoff activity [27]. Nevertheless, all the amino acid restudies identified in this study were located at the N-terminal region (**Figure 2**). We mapped the identified amino acids in the N-terminal structure of PA. Interestingly, amino acid mutations identified in our screening were widely distributed in the N-terminal region of 1918 PA-X (**Figure 8**).

**Figure 8:**
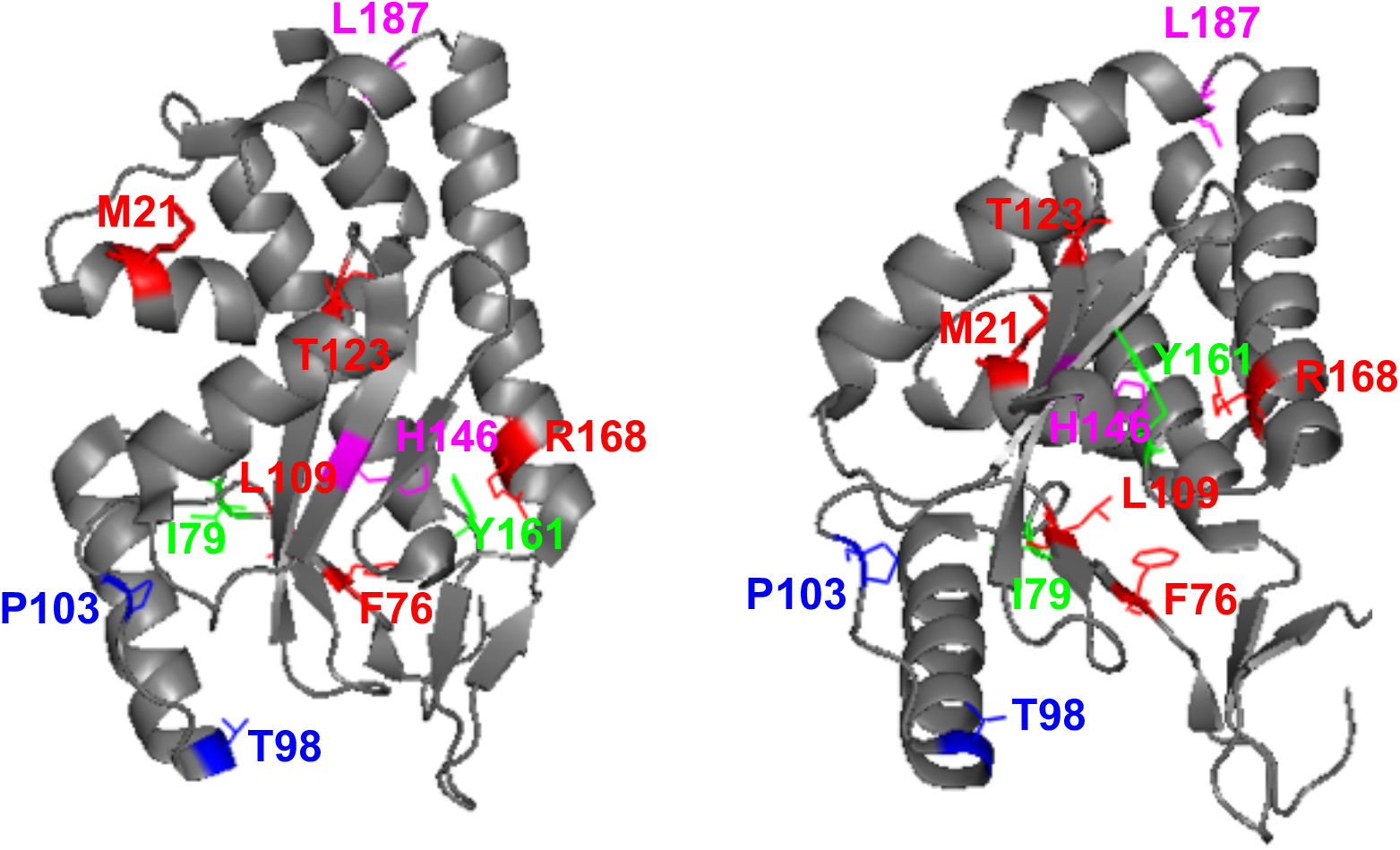
Location of identified amino acid mutations in A/California/04/2009-pH1N1. The N-terminal region of pH1N1 PA protein (Accession #AY818132) is shown in two different views using the same coloring code. PA is indicated in gray. Location of amino acids M21, F76, L109, T123, R168 (red), T98/P103 (blue), I79/Y161 (green) and H146/L187 (magenta) are shown on the background of the PA. The crystal structure figure was adapted from PDB: 4AVQ using Pymol.

Amino acid changes affecting 1918 PA-X’s shutoff activity also affected the ability of 1918 PA-X to inhibit IFN responses after SeV infection measured by the expression levels of Fluc driven by and ISRE element (**Figure 7A**), as previously shown for other IAV PA-X mutants [34]. Given that IFN responses include the induction of genes with antiviral activity, controlling viral replication [39], and that PA-X modulates virus pathogenesis [18], it is very likely that the identified mutations in PA-X affect virus fitness and pathogenesis.

Remarkably, amino acid residues present in 1918 PA-X WT are highly conserved (> 99.7%) in human H1N1 viruses isolated since 1918, including 2009 pH1N1 viruses (**Table 2**). These results strongly suggest that H1N1 IAV encoding PA-X proteins affected in inhibition of host shutoff present disadvantages for replication in humans, and therefore, they are not present in nature. Alternatively, given that IAV PA-X shares its N-terminal 191 amino acids with PA [9], it could be that these residues are important for PA’s polymerase activity, for PA-X shutoff activity, or both. In line with this hypothesis, it has been shown that residues P103, and L109 are important for PA activity [40,41].

The relevance for PA-X shutoff activity of three of the amino acid residues identified in our screening (T123, H146 and R168) were recently reported in another manuscript in which authors used a yeast-based assay to identify amino acid changes involved in WSN H1N1 PA-X inhibition of host gene expression [28], further validating our results. Whereas the amino acid change at position 146 (H146Y) was exactly the same in our work and in the reported manuscript, for positions 123 and 168 the amino acid changes were different: T123I and R168M in [28] and T123A and R168G in our work. Moreover, these results suggest that the contribution of these amino acid residues to inhibit host gene expression are not strain specific. In the case of the PA-X I79V/Y161, T98I/P103S, and H146Y/L187P double mutants, we determined whether one or both amino acid residues were required for 1918 PA-X shutoff activity. Our results indicate that all the amino acid changes contributed to decrease 1918 PA-X’s ability to induce cellular shutoff (**Figures 5 and 6**). Similarly, in a previous work using an H5N1 IAV strain, we also found that amino acid mutations H146Y, and L187P, present in the double mutants, as well as F76L, found in a single mutant, decrease the ability of H5N1 PA-X to inhibit host gene expression [36].

In this study we validate a convenient and novel approach, based in the simply use of bacteria, for the identification of residues relevant for IAV PA-X’s inhibition of host gene expression. This system has allowed us to identify 1918 PA-X amino acid residues important for its shutoff activity and for the ability of the protein to impair IFN responses after viral infection, which likely has implications in virus pathogenesis. Interestingly, the importance of some of the identified 1918 PA-X residues for host shutoff activity were previously shown with other IAV strains, further validating the use of this bacteria-based approach to identify mutations affecting IAV PA-X’s inhibition of cellular host gene expression. These results demonstrate the feasibility of using this bacteria-based system to identify other viral proteins inhibiting host gene expression as well as amino acid residues involved in host shutoff of these viral proteins. Because functional relevance and high conservation of these amino acids, it could be feasible to design novel compounds targeting these protein regions for the treatment of influenza infections.

## Author Contributions

Conceptualization, K.C., L.M-S., A.N. and M.L.D.; formal analysis, K.C., L.M-S., A.N. and M.L.D.; investigation, K.C., L.M-S., A.N. and M.L.D.; data curation, K.C., L.M-S., A.N. and M.L.D.; writing—original draft preparation, K.C., and M.L.D.; writing—review and editing, K.C., L.M-S., A.N. and M.L.D.; funding acquisition, L.M-S., A.N. and M.L.D. All authors have read and agreed to the published version of the manuscript.

## Institutional Review Board Statement

Not applicable.

## Informed Consent Statement

Not applicable.

## Data Availability Statement

The data that support the findings of this study are available from the corresponding author upon reasonable request.

## Acknowledgments

We would like to thank members at our institutes for their efforts in keeping them fully operational during the COVID-19 pandemic.

## References

1. Cheung, T.K.W., Poon, L.L.M. Biology of Influenza a Virus. Ann N Y Acad Sci 2007, 1102, 1–25, doi:10.1196/annals.1408.001.

2. Wille, M., Holmes, E.C. The Ecology and Evolution of Influenza Viruses. Cold Spring Harb Perspect Med 2020, 10, doi:10.1101/cshperspect.a038489.

3. Tong, S., Zhu, X., Li, Y., Shi, M., Zhang, J., Bourgeois, M., Yang, H., Chen, X., Recuenco, S., Gomez, J., et al. New World Bats Harbor Diverse Influenza A Viruses. PLoS Pathog 2013, 9, e1003657, doi:10.1371/journal.ppat.1003657.

4. Nogales, A., Chauché, C., DeDiego, M.L., Topham, D.J., Parrish, C.R., Murcia, P.R., Martínez-Sobrido, L. The K186E Amino Acid Substitution in the Canine Influenza Virus H3N8 NS1 Protein Restores Its Ability To Inhibit Host Gene Expression. J Virol 2017, 91, doi:10.1128/JVI.00877-17.

5. Parrish, C.R., Murcia, P.R., Holmes, E.C. Influenza Virus Reservoirs and Intermediate Hosts: Dogs, Horses, and New Possibilities for Influenza Virus Exposure of Humans. J Virol 2015, 89, 2990–2994, doi:10.1128/JVI.03146-14.

6. Taubenberger, J.K., Morens, D.M. Pandemic Influenza--Including a Risk Assessment of H5N1. Rev Sci Tech 2009, 28, 187–202, doi:10.20506/rst.28.1.1879.

7. Taubenberger, J.K., Morens, D.M. 1918 Influenza: The Mother of All Pandemics. Emerg Infect Dis 2006, 12, 15–22, doi:10.3201/eid1201.050979.

8. Johnson, N.P.A.S., Mueller, J. Updating the Accounts: Global Mortality of the 1918-1920 “Spanish” Influenza Pandemic. Bull Hist Med 2002, 76, 105–115, doi:10.1353/bhm.2002.0022.

9. Jagger, B.W., Wise, H.M., Kash, J.C., Walters, K.-A., Wills, N.M., Xiao, Y.-L., Dunfee, R.L., Schwartzman, L.M., Ozinsky, A., Bell, G.L., et al. An Overlapping Protein-Coding Region in Influenza A Virus Segment 3 Modulates the Host Response. Science 2012, 337, 199–204, doi:10.1126/science.1222213.

10. Yuan, P., Bartlam, M., Lou, Z., Chen, S., Zhou, J., He, X., Lv, Z., Ge, R., Li, X., Deng, T., et al. Crystal Structure of an Avian Influenza Polymerase PA(N) Reveals an Endonuclease Active Site. Nature 2009, 458, 909–913, doi:10.1038/nature07720.

11. Dias, A., Bouvier, D., Crépin, T., McCarthy, A.A., Hart, D.J., Baudin, F., Cusack, S., Ruigrok, R.W.H. The Cap-Snatching Endonuclease of Influenza Virus Polymerase Resides in the PA Subunit. Nature 2009, 458, 914–918, doi:10.1038/nature07745.

12. Khaperskyy, D.A., Schmaling, S., Larkins-Ford, J., McCormick, C., Gaglia, M.M. Selective Degradation of Host RNA Polymerase II Transcripts by Influenza A Virus PA-X Host Shutoff Protein. PLoS Pathog 2016, 12, e1005427, doi:10.1371/journal.ppat.1005427.

13. Gao, H., Xu, G., Sun, Y., Qi, L., Wang, J., Kong, W., Sun, H., Pu, J., Chang, K.-C., Liu, J. PA-X Is a Virulence Factor in Avian H9N2 Influenza Virus. J Gen Virol 2015, 96, 2587–2594, doi:10.1099/jgv.0.000232.

14. Desmet, E.A., Bussey, K.A., Stone, R., Takimoto, T. Identification of the N-Terminal Domain of the Influenza Virus PA Responsible for the Suppression of Host Protein Synthesis. J Virol 2013, 87, 3108–3118, doi:10.1128/JVI.02826-12.

15. Khaperskyy, D.A., Emara, M.M., Johnston, B.P., Anderson, P., Hatchette, T.F., McCormick, C. Influenza a Virus Host Shutoff Disables Antiviral Stress-Induced Translation Arrest. PLoS Pathog 2014, 10, e1004217, doi:10.1371/journal.ppat.1004217.

16. Wang, X.-H., Gong, X.-Q., Wen, F., Ruan, B.-Y., Yu, L.-X., Liu, X.-M., Wang, Q., Wang, S.-Y., Wang, J., Zhang, Y.-F., et al. The Role of PA-X C-Terminal 20 Residues of Classical Swine Influenza Virus in Its Replication and Pathogenicity. Vet Microbiol 2020, 251, 108916, doi:10.1016/j.vetmic.2020.108916.

17. Liu, L., Song, S., Shen, Y., Ma, C., Wang, T., Tong, Q., Sun, H., Pu, J., Iqbal, M., Liu, J., et al. Truncation of PA-X Contributes to Virulence and Transmission of H3N8 and H3N2 Canine Influenza Viruses in Dogs. J Virol 2020, 94, doi:10.1128/JVI.00949-20.

18. Nogales, A., Martinez-Sobrido, L., Topham, D.J., DeDiego, M.L. Modulation of Innate Immune Responses by the Influenza A NS1 and PA-X Proteins. Viruses 2018, 10, doi:10.3390/v10120708.

19. Hu, J., Mo, Y., Gao, Z., Wang, X., Gu, M., Liang, Y., Cheng, X., Hu, S., Liu, W., Liu, H., et al. PA-X-Associated Early Alleviation of the Acute Lung Injury Contributes to the Attenuation of a Highly Pathogenic H5N1 Avian Influenza Virus in Mice. Med Microbiol Immunol 2016, 205, 381–395, doi:10.1007/s00430-016-0461-2.

20. Gao, H., Sun, Y., Hu, J., Qi, L., Wang, J., Xiong, X., Wang, Y., He, Q., Lin, Y., Kong, W., et al. The Contribution of PA-X to the Virulence of Pandemic 2009 H1N1 and Highly Pathogenic H5N1 Avian Influenza Viruses. Sci Rep 2015, 5, 8262, doi:10.1038/srep08262.

21. Hu, J., Mo, Y., Wang, X., Gu, M., Hu, Z., Zhong, L., Wu, Q., Hao, X., Hu, S., Liu, W., et al. PA-X Decreases the Pathogenicity of Highly Pathogenic H5N1 Influenza A Virus in Avian Species by Inhibiting Virus Replication and Host Response. J Virol 2015, 89, 4126–4142, doi:10.1128/JVI.02132-14.

22. Gao, H., Xu, G., Sun, Y., Qi, L., Wang, J., Kong, W., Sun, H., Pu, J., Chang, K.-C., Liu, J. PA-X Is a Virulence Factor in Avian H9N2 Influenza Virus. J Gen Virol 2015, 96, 2587–2594, doi:10.1099/jgv.0.000232.

23. Lee, J., Yu, H., Li, Y., Ma, J., Lang, Y., Duff, M., Henningson, J., Liu, Q., Li, Y., Nagy, A., et al. Impacts of Different Expressions of PA-X Protein on 2009 Pandemic H1N1 Virus Replication, Pathogenicity and Host Immune Responses. Virology 2017, 504, 25–35, doi:10.1016/j.virol.2017.01.015.

24. Gao, H., Sun, H., Hu, J., Qi, L., Wang, J., Xiong, X., Wang, Y., He, Q., Lin, Y., Kong, W., et al. Twenty Amino Acids at the C-Terminus of PA-X Are Associated with Increased Influenza A Virus Replication and Pathogenicity. J Gen Virol 2015, 96, 2036– 2049, doi:10.1099/vir.0.000143.

25. Hayashi, T., Chaimayo, C., McGuinness, J., Takimoto, T. Critical Role of the PA-X C-Terminal Domain of Influenza A Virus in Its Subcellular Localization and Shutoff Activity. J Virol 2016, 90, 7131–7141, doi:10.1128/JVI.00954-16.

26. Sun, Y., Hu, Z., Zhang, X., Chen, M., Wang, Z., Xu, G., Bi, Y., Tong, Q., Wang, M., Sun, H., et al. An R195K Mutation in the PA-X Protein Increases the Virulence and Transmission of Influenza A Virus in Mammalian Hosts. J Virol 2020, 94, doi:10.1128/JVI.01817-19.

27. Oishi, K., Yamayoshi, S., Kawaoka, Y. Mapping of a Region of the PA-X Protein of Influenza A Virus That Is Important for Its Shutoff Activity. J Virol 2015, 89, 8661–8665, doi:10.1128/JVI.01132-15.

28. Oishi, K., Yamayoshi, S., Kawaoka, Y. Identification of Novel Amino Acid Residues of Influenza Virus PA-X That Are Important for PA-X Shutoff Activity by Using Yeast. Virology 2018, 516, 71–75, doi:10.1016/j.virol.2018.01.004.

29. Kochs, G., García-Sastre, A., Martínez-Sobrido, L. Multiple Anti-Interferon Actions of the Influenza A Virus NS1 Protein. J Virol 2007, 81, 7011–7021, doi:10.1128/JVI.02581-06.

30. Rosano, G.L., Ceccarelli, E.A. Recombinant Protein Expression in Escherichia Coli: Advances and Challenges. Front Microbiol 2014, 5, 172, doi:10.3389/fmicb.2014.00172.

31. Levene, R.E., Shrestha, S.D., Gaglia, M.M. The Influenza A Virus Host Shutoff Factor PA-X Is Rapidly Turned over in a Strain-Specific Manner. J Virol 2021, doi:10.1128/JVI.02312-20.

32. Nogales, A., Martinez-Sobrido, L., Topham, D.J., DeDiego, M.L. Modulation of Innate Immune Responses by the Influenza A NS1 and PA-X Proteins. Viruses 2018, 10, doi:10.3390/v10120708.

33. Nogales, A., Rodriguez, L., DeDiego, M.L., Topham, D.J., Martínez-Sobrido, L. Interplay of PA-X and NS1 Proteins in Replication and Pathogenesis of a Temperature-Sensitive 2009 Pandemic H1N1 Influenza A Virus. J Virol 2017, 91, doi:10.1128/JVI.00720-17.

34. Nogales, A., Martinez-Sobrido, L., Chiem, K., Topham, D.J., DeDiego, M.L. Functional Evolution of the 2009 Pandemic H1N1 Influenza Virus NS1 and PA in Humans. J Virol 2018, 92, doi:10.1128/JVI.01206-18.

35. Gaucherand, L., Porter, B.K., Levene, R.E., Price, E.L., Schmaling, S.K., Rycroft, C.H., Kevorkian, Y., McCormick, C., Khaperskyy, D.A., Gaglia, M.M. The Influenza A Virus Endoribonuclease PA-X Usurps Host MRNA Processing Machinery to Limit Host Gene Expression. Cell Rep 2019, 27, 776-792.e7, doi:10.1016/j.celrep.2019.03.063.

36. Chiem, K., López-García, D., Ortego, J., Martinez-Sobrido, L., DeDiego, M.L., Nogales, A. Identification of Amino Acid Residues Required for Inhibition of Host Gene Expression by Influenza A/Viet Nam/1203/2004 H5N1 PA-X. J Virol 2021, doi:10.1128/JVI.00408-21.

37. Shi, M., Jagger, B.W., Wise, H.M., Digard, P., Holmes, E.C., Taubenberger, J.K. Evolutionary Conservation of the PA-X Open Reading Frame in Segment 3 of Influenza A Virus. J Virol 2012, 86, 12411–12413, doi:10.1128/JVI.01677-12.

38. Chen, X., Wang, W., Wang, Y., Lai, S., Yang, J., Cowling, B.J., Horby, P.W., Uyeki, T.M., Yu, H. Serological Evidence of Human Infections with Highly Pathogenic Avian Influenza A(H5N1) Virus: A Systematic Review and Meta-Analysis. BMC Med 2020, 18, 377, doi:10.1186/s12916-020-01836-y.

39. Iwasaki, A., Pillai, P.S. Innate Immunity to Influenza Virus Infection. Nat Rev Immunol 2014, 14, 315–328, doi:10.1038/nri3665.

40. Hara, K., Schmidt, F.I., Crow, M., Brownlee, G.G. Amino Acid Residues in the N-Terminal Region of the PA Subunit of Influenza A Virus RNA Polymerase Play a Critical Role in Protein Stability, Endonuclease Activity, Cap Binding, and Virion RNA Promoter Binding. J Virol 2006, 80, 7789–7798, doi:10.1128/JVI.00600-06.

41. Zhao, H., Chu, H., Zhao, X., Shuai, H., Wong, B.H.-Y., Wen, L., Yuan, S., Zheng, B.-J., Zhou, J., Yuen, K.-Y. Novel Residues in the PA Protein of Avian Influenza H7N7 Virus Affect Virulence in Mammalian Hosts. Virology 2016, 498, 1–8, doi:10.1016/j.virol.2016.08.004.

